# Root associated bacterial communities and root metabolite composition are linked to nitrogen use efficiency in sorghum

**DOI:** 10.1101/2023.02.08.527764

**Authors:** Yen Ning Chai, Yunhui Qi, Emily Goren, Amy M. Sheflin, Susannah Tringe, Jessica E. Prenni, Peng Liu, Daniel Schachtman

## Abstract

Development of cereal crops with high nitrogen-use efficiency (NUE) is a priority for worldwide agriculture. In addition to conventional plant breeding and genetic engineering, the use of the plant microbiome offers another approach to improve crop NUE. To gain insight into the bacterial communities associated with sorghum lines that differ in NUE, a field experiment was designed comparing 24 diverse sorghum lines under sufficient and deficient nitrogen (N). Amplicon sequencing and untargeted gas chromatography-mass spectrometry (GC-MS) were used to characterize the bacterial communities and the root metabolome associated with sorghum genotypes varying in sensitivity to low N. We demonstrated that N stress and sorghum type (energy, sweet, and grain sorghum) significantly influenced the root-associated bacterial communities and root metabolite composition of sorghum. Sorghum NUE was positively correlated with the bacterial richness and diversity in the rhizosphere. The greater alpha diversity in high NUE lines was associated with the decreased abundance of a dominant bacterial taxa, *Pseudomonas*. Multiple strong correlations were detected between root metabolites and rhizosphere bacterial communities in response to N stress and indicate that the shift in the sorghum microbiome due to low-N is associated with the root metabolites of the host plant. Taken together, our study provides new insight into the links between host genetic regulation of root metabolites and root-associated microbiome of sorghum genotypes differing in NUE and tolerance to low-N stress.

## Introduction

The association between root metabolites and the microbes that inhabit both the inside of the roots (endosphere) and in the region just outside the roots (rhizosphere) are integral parts of plant microbe interactions. Although it is known that certain plant metabolites exuded from plant roots can modulate the assembly of the rhizosphere microbiome (Lundberg et al., 2012; Pascale et al., 2020; Stringlis et al., 2018; Zhalnina et al., 2018), the effect from a substantial number of primary and secondary root metabolites on the root-associated microbiome remains elusive. Few studies have characterized the association between the root metabolites and the root-associated microbiome (Huang et al., 2019; Lebeis et al., 2015; Stringlis et al., 2018) and even fewer studies have investigated how this association is affected by abiotic stresses (Caddell et al., 2020; Sheflin et al., 2019). Since changes in root metabolites are an important phenotypic trait that could occur as a result of alterations in environmental conditions (Bino et al., 2004; Fiehn, 2002; Roessner et al., 2001), they will also provide additional clues as to the factors that underlie the subtle host plant modulation of root-associated microbiome to overcome the stress. In addition, the interaction between root exudates and root-associated microbiome is likely to depend on plant species and genotype being considered. Host plants have been shown to have significant effect on the composition of root-associated microbiome (Fitzpatrick et al., 2018; Lebeis et al., 2015; Liu et al., 2019; Peiffer et al., 2013). Understanding the host plant control over root-associated microbiome is also of great interest to crop breeders to select for cultivars that are more productive and resilient to stresses.

Over the past few decades, the application of nitrogen fertilizer has intensified to accommodate increased demands for higher crop yields due to rapidly growing populations. While excess amount of applied N ensures higher yields, it also leads to adverse environmental impacts due to the leaching of N fertilizer into waterways and groundwater. Therefore, to mitigate adverse environmental effects it will be necessary to lower the input of N fertilizer while selecting for crop cultivars with high N use efficiency. Sorghum is an excellent candidate for studying this because it requires less N fertilizer (Moore et al., 2021) and exhibits a range of nitrogen use efficiency (NUE) which may be in part due to the vast genetic diversity within the sorghum gene pool.While past studies have identified genes (Bollam et al., 2021; Gelli et al., 2014) and phenotypes (Borrell et al., 2001; Gardner et al., 1994) that are potentially associated with high NUE in sorghum, there is still a lack of information on the overall variation in sorghum NUE and the relationship between NUE and recruitment of root-associated microbes.

As an ancient African grass, elite sorghum lines that are grown as a staple crop worldwide are the result of many generations of breeding concomitant with the selection across geographic gradients (Boatwright et al., 2021). Sorghum can be classified into four major types based on their carbon partitioning characteristics (Boatwright et al., 2021). Grain sorghum is the most widely grown sorghum type, is primarily grown for food and partitions carbon to panicles. The forage cultivars have lower grain yield and exhibit coarser stems (as compared to grain sorghum) and are used mainly for grazing and silage. In contrast, energy and sweet sorghum mostly partition carbon to the stem but in the form of lignocellulosic biomass and fermentable sugars, respectively, making them excellent feedstocks for biofuel production (Mathur et al., 2017; Mullet et al., 2014). Information about the NUE of energy sorghum will be particularly important to identify lines that grow well in marginal soils with low-N to increase cellulosic biomass production in a low input system.

In this study, we characterized the root metabolite and bacterial communities across 24 diverse sorghum genotypes grown under full- and low-N field conditions. Our goal was to construct a more comprehensive picture that depicts the interplay between the host regulation of root metabolites and root-associated microbiome of sorghum genotypes differing in response to N stress. To achieve this goal, we sought to answer the following questions: (*i*) How does N stress and sorghum genotype affect the associated bacterial communities?, (*ii*) How do N stress and sorghum genotype affect the root metabolite composition?, (*iii*) Are the bacterial communities different between high and low NUE genotypes?, (*iv*) Is there an association between root metabolites and rhizosphere bacterial community composition? This is vital foundational information for our overall understanding of the relationship between root metabolites and associated microbes and could be used in future work to intentionally promote specific plant microbe interactions that would be more favorable to low N conditions for sorghum and for other crop species.

## Materials and Methods

### Field description and experimental design

The field used in this study was located in Central Nebraska, USA at GPS coordinates 41.201041, −97.944750. Pre-season soil nitrogen was measured and found to be very low (0.6 ppm nitrate-N). The field was set up as a split plot design with eight blocks or replicates that contained 2 treatments and 24 sorghum genotypes (Table 1) arranged randomly within each treatment. Eighty-five pounds of granular urea (46-0-0) was added to the full-N treatments and no urea was added to the low-N treatments. Seeds were treated with Concep III before planting and metolachlor and bromoxynil were used just after sorghum germination to help control weeds. Planting was performed on May 26, 2017 and seeds were planted at an interval of 4 inches in a row in four row plots that were 10 foot in length. Mechanical cultivation was done by hand labor during the season.

**Table 1.** Sorghum genotypes used in this study and the types of data collected for each genotype. The collection of samples for microbiome characterization was performed on all genotypes. Biomass data was measured for all genotypes but Chinese Amber due to severe lodging prior to the harvest.

| Genotype | Type | Race | Root Metabolite |
| --- | --- | --- | --- |
| PI 329311 | Energy | Durra | Yes |
| PI 297155 |  | Kafir | Yes |
| PI 506069 |  | Guinea/bicolor | Yes |
| PI 297130 |  | Caudatum | Yes |
| Grassl |  | Caudatum | No |
| PI 152730 |  | Caudatum/bicolor | Yes |
| PI 655972 |  | Kafir | No |
| PI 510757 |  | Durra | Yes |
| PI 505735 |  | Caudatum | No |
| PI 329632 |  | Durra | Yes |
| PI 35038 |  | Caudatum | Yes |
| PI 585954 |  | Guinea | Yes |
| NTJ2 |  | Durra | Yes |
| M81e |  | Caudatum/durra | Yes |
| PI 229841 |  | Kafir | Yes |
| BTx623 | Grain | Kafir | No |
| Btx642 |  | Caudatum | Yes |
| China 17 | Sweet | Bicolor | Yes |
| San Chi San |  | Bicolor | Yes |
| ICSV700 |  | Bicolor | Yes |
| Atlas |  | Kafir | Yes |
| Leoti |  | Kafir/bicolor | Yes |
| Chinese Amber |  | Bicolor | No |
| Rio |  | Durra/caudatum | Yes |

### Sample collection and preparation

#### Biomass sampling

Final plant biomass was measured on October 9, 2017 by harvesting a 1 meter section of one of the middle rows and then using the area of that section to extrapolate to kg/hectare. Stalks and panicles were weighed separately. Four replicate blocks out of the eight were subsampled for dry weight. Plants were weighed and a subsample of three plants was taken then reweighed, bagged, dried at 80 °C and reweighed to determine the dry weights of the plots. We computed a biomass ratio by dividing the total aboveground biomass under low-by the biomass under full-N for each genotype. This ratio was used to evaluate the nitrogen-use efficiency (NUE) of the different sorghum genotypes under N limited conditions. The biomass data of one genotype Chinese Amber could not be measured due to severe lodging prior to the biomass harvest.

#### Microbiome and root metabolite sampling

Soil between rows, soil within rows, rhizosphere, root, and leaf samples were collected from the field on July 18, 2017. Except for the soil between rows, each sample was collected from two plants located at two different spots within each plot. Soil between rows samples were excavated from the top 30 cm of soil in between plots from different locations of the field. Soil within rows, rhizosphere, and root samples were collected as described previously (McPherson et al., 2018). Leaf samples were collected from the newest, unexpanded leaves inside the whorl that had never been exposed because they were considered to be absent of leaf epiphytic microbes. The leaf tissue was rinsed in phosphate buffer and cut into small pieces by handling carefully with sterile tools and then stored on ice. In addition to collecting root samples for microbiome analysis, a subset of roots was separated for metabolite profiling after carefully removing the rhizosphere soil by rinsing again in phosphate buffer and gently wiping the roots with a Kimwipe.

All the samples for microbiome and metabolome analyses were brought back to the laboratory on ice and processed as described previously (McPherson et al., 2018; Sheflin et al., 2019). The soil between rows were processed with the same procedure as the soil within rows. In brief, these soil samples were sieved to remove any small roots and debris before freezing at −20 °C and then loading into 96-well plate for DNA extraction. For the leaf samples, a surface sterilization step was omitted because the leaf portion that we collected was still inside the plant and had not emerged from the shoot. Both root tissues and leaf samples were frozen at −80 °C and ground in liquid N to homogenize the samples before DNA extraction.

### DNA extraction and sequencing

Following the manufacturer’s protocols, soil and rhizosphere were extracted using MagAttract PowerSoil DNA KF Kit (Qiagen, Germantown, MD) with KingFisher Flex System (ThermoFisher Scientific, Waltham, MA) while MagMAX™ Plant DNA Kit (ThermoFisher Scientific, Waltham, MA) was used for the extraction of leaf and root tissues. The extracted 16S rRNA was then amplified at the V4 region with the primers F515/R806 followed by sequencing with Illumina MiSeq platform at the Joint Genome Institute using the protocol described previously (Chiniquy et al., 2021).

### Non-targeted GC-MS and data analysis

Non-targeted gas chromatography-mass spectrometry (GC-MS) was used for root metabolite profiling using the protocol described previously (Sheflin et al., 2019). Root metabolite profiling was performed on 19 selected genotypes (excluded Grassl, PI655972, PI505735, BTx623, and Chinese Amber) with five replicates for each genotype and treatment combination. Principal component analysis was performed on all metabolites (annotated metabolites and unknowns) to visualize the overall variation in the data. The relative intensity of the annotated metabolites was clustered by mean using one-dimensional self-organizing map (1D-SOM) (Meinicke et al., 2008). The 1D-SOM classified the metabolites with similar intensity patterns into ten clusters for the high- and low-NUE lines under both N treatments.

### 16S amplicon data analysis

The analysis of 16S amplicon data was done using UPARSE (Edgar, 2013) and QIIME 2 (Bolyen et al., 2019) and R was used for figure generation. UPARSE was used to merge the paired-end reads, remove the primers, filter out the reads with expected error scores below one, remove chimeras, and generate read clusters at 100 % similarity.

The resulting table containing the read count of the amplicon sequence variants (ASV) was then subjected to downstream analyses using QIIME 2. Taxonomy was assigned to each ASV using a q2-feature-classifier (Bokulich et al., 2018) pre-trained with SILVA database (Quast et al., 2013). Rarefaction curves were generated for each sample type prior to the alpha and beta diversity analyses to ensure sufficient sampling depth was achieved. Sampling depth of 84009, 57118, 80141, 15496, and 500 were used for soil between rows, soil within rows, rhizosphere, root endosphere, and leaf, respectively.For the alpha diversity analyses, Shannon diversity index and Faith’s phylogenetic distance were used to access the diversity and richness of the microbiome, respectively. The difference in alpha diversity indices for each combination of N treatment and sample type was tested using ANOVA in R followed by *post-hoc* Tukey-HSD pairwise comparisons. For the beta diversity analyses, principal coordinate analysis (PCoA) and constrained analysis of principal coordinates (CAP) were performed on Bray-Curtis distance matrices using *pcoa* (Paradis E., 2019) and *capscale* (Oksanen et al., 2015) functions in R. The changes in microbiome composition due to N treatment, sample type, sorghum genotype, sorghum type (grain, energy, sweet) were evaluated using PERMANOVA with 999 permutations using *adonis* function (Oksanen et al., 2015) in R. The bacterial taxa differentially affected by N treatment and sorghum NUE were identified using linear discriminant analysis (LDA) effect size (LEfSe) analysis at http://huttenhower.sph.harvard.edu/lefse/ (Segata et al., 2011).

### Correlation between metabolites and rhizosphere microbiome

Group sparse canonical correlation analysis (GSCCA) was used to identify the correlations between root metabolites and rhizosphere ASVs (Lin et al., 2013). Only the samples that were profiled for both root metabolite and microbiome were included in this analysis. The ASVs present in less than 95 % of the samples for any combination of sorghum type and genotype in both low- and full-N treatments were excluded from the analysis. Metabolites that were unannotated were also excluded from GSCCA. As a result, a total of 690 ASVs and 77 metabolites were included in the GSCCA. The filtered ASV data was subjected to ANCOM (analysis of composition of microbiomes) transformation (Kaul et al., 2017) while the metabolite data was log-transformed. Next, we grouped the ASVs and metabolites based on their phylum and superclass, respectively. Superclass is the second-level chemical taxonomy assigned to biochemicals according to their structural features (The Metabolomics Workbench, https://www.metabolomicsworkbench.org/). We obtained the between group and within group correlations and the groups with high between group correlations were merged until the within group correlations were higher than the between group correlations. We also merged the superclass or phylum singletons with the corresponding most correlated groups. After merging, we performed GSCCA based on the pooled covariance with respect to full- and low-N groups to select groups and features that contribute to the canonical correlations between metabolites and ASVs, followed by a test with 10,000 permutations to determine if the canonical correlations were significantly different from zero. To account for the imbalance of sample sizes in different genotypes, we used stratified cross validation when searching for the best tuning parameters.

### Data availability

The 16S rRNA sequences used in this study are available at the NCBI repository under the BioProject number PRJNA830495.

## Results

### Diverse panel of sorghum genotypes exhibit variation in N-use efficiency

In this study, the NUE of 23 diverse sorghum genotypes that span three sorghum types were compared (Figure 1, Figure S1). The sorghum lines were grown under full- and low-N conditions in the same field site and their NUE was derived as the ratio of the aboveground biomass under low-versus full-N conditions measured at the end of the season. The low-N treatment significantly reduced the biomass of all the sorghum genotypes (ANOVA: *P* < 0.001). The average reduction in dry biomass due to low-N was 22 % with some genotypes exhibiting larger reductions in biomass than others. PI 329632 was the most susceptible to low-N, exhibiting 40 % reduction in dry biomass in response to low-N. ICSV700 was the least susceptible to low-N as its biomass barely decreased due to low N. The sweet sorghum genotypes used in this study exhibited an average of 23 % higher dry biomass ratio than the other two sorghum types, indicating higher NUE in this sorghum type.

**Figure 1.**
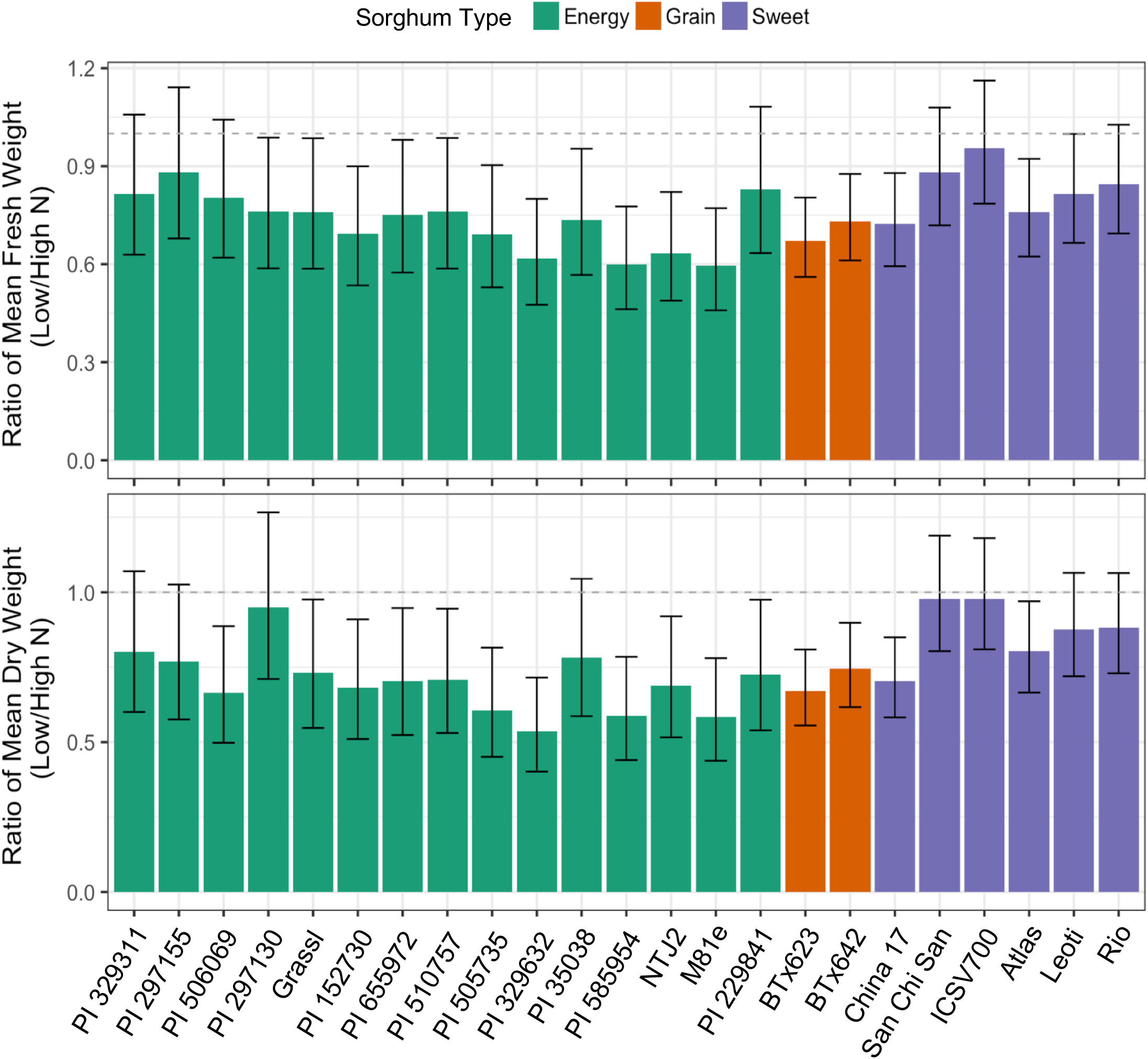
Biomass ratios of 23 diverse sorghum genotypes. Fresh (top) and dry (bottom) weight ratio derived by dividing the biomass measured under low-*vs.* full-N conditions. Error bars represent the 95 % confidence intervals.

### Sorghum rhizosphere and root endosphere microbiome are affected by N availability and sorghum type

The bacterial communities in the soil between rows, soil within rows (the soil layer beyond rhizosphere but near roots), rhizosphere, root endosphere, and leaf endosphere at sorghum vegetative stage were surveyed by sequencing the V4 region of the 16S rRNA amplicons. We found that sample type or compartment (rhizosphere, root and leaf endosphere, soils) were the major factors that affected the assembly of sorghum microbiome (Figure 2A), accounting for 27 % of the variation in the microbial communities. The microbial diversity and richness decreased from the soil between rows to the endophytic compartments, with the leaf endosphere harboring the least diverse microbial community (Figure 2B,C). No significant change in these alpha diversity indices was detected in these compartments due to low-N (Figure 2B,C). From the soil to the endophytic compartments, we observed a decrease in relative abundance of the members from the phylum Actinobacteriota, Firmicutes, Crenarchaeota, and Chloroflexi, while an increase in abundance was found for Bacteroidota and Proteobacteria (Figure 2D).

**Figure 2.**
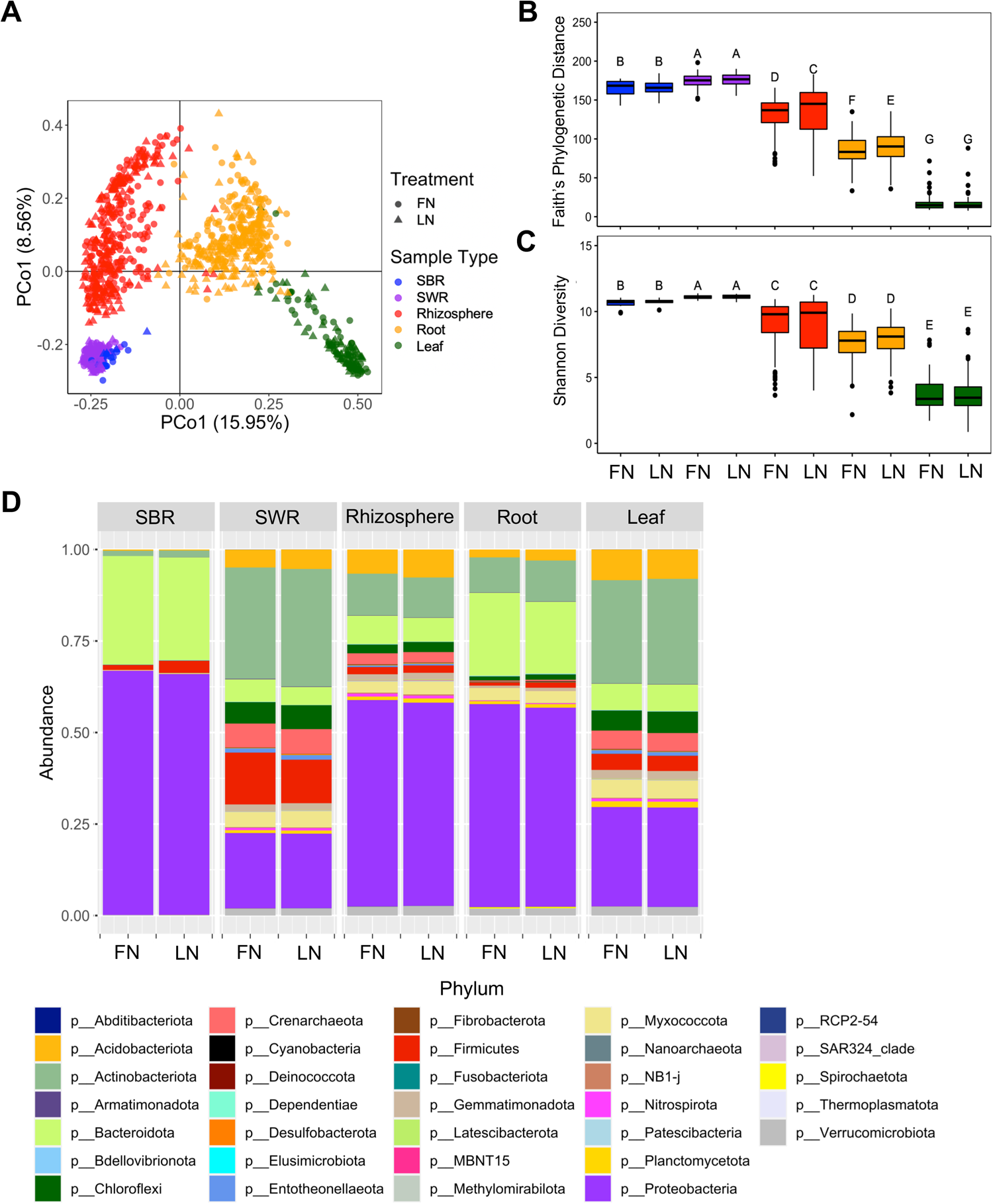
Bacterial diversity and composition across different compartments. (A) PCoA based on Bray Curtis distance showing the overall composition of bacterial communities across different compartments and N treatments. (B) Faith’s phylogenetic distance and (C) Shannon diversity of bacterial communities for each compartment and treatment combination. (D) Relative abundance of the most abundant phyla across different compartments and N treatments. Abbreviation: SWR: soil within rows; SBR: soil between rows; FN: full-N; LN: low-N.

Next, we dissected the effects of N availability and sorghum type on the bacterial communities for each compartment (Figure 3A-E). Canonical analysis of principal coordinates (CAP) and PERMANOVA revealed that the rhizosphere and root endosphere communities were significantly impacted by N availability, which separated the samples along the primary axis (Figure 3B,C), suggesting that it was a major factor that shaped the microbial communities in these compartments. Linear discriminant analysis effect size (LEfSE) was then used to identify the microbial taxa across all taxonomic ranks that were differentially impacted by N availability for each compartment. Consistent with the CAP and PERMANOVA findings, significant differential abundance at the phyla level were only detectable in the rhizosphere and root endosphere (Figure 4). Bacteroidota was the only phylum that was significantly enriched under full-N conditions and this was observed for both rhizosphere and root endosphere. Chloroflexi and Planctomycetota were in greater abundance under low-N condition in both rhizosphere and endosphere, while Myxococcota was only enriched in low-N rhizosphere samples. The phyla that were significantly enriched under low-N conditions in the root endosphere were Firmicutes, Acidobacteria, Crenarachaeota, and Gemmatimonadota. In addition to N availability, we observed a significant effect from sorghum type (energy, grain and sweet) in all compartments except for soil between rows which was the only compartment that was not associated with plants (Figure 3A-E). Subsequent pairwise PERMANOVA revealed that the microbial communities were more similar between the sweet and energy types, as compared with the grain sorghum (q < 0.05).

**Figure 3.**
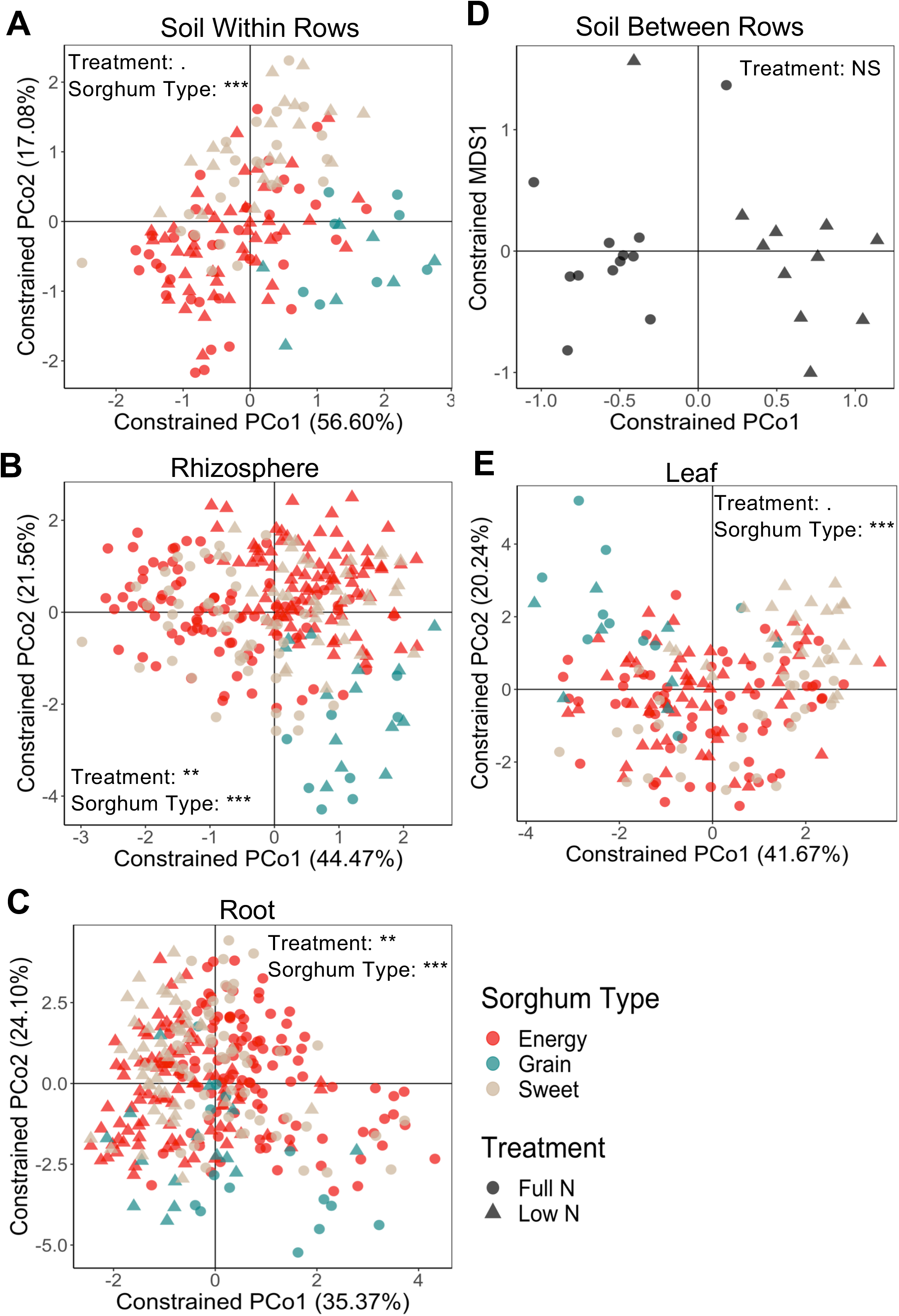
Effects of N treatment and sorghum type on the bacterial community in each compartment. CAP based on Bray Curtis distance of (A) soil within rows, (B) rhizosphere, (C) root endosphere, (D) soil between rows, and (E) leaf endosphere. *p ≤ 0.05, **p ≤ 0.01, ***p ≤ 0.001. Abbreviation: NS: not significant.

**Figure 4.**
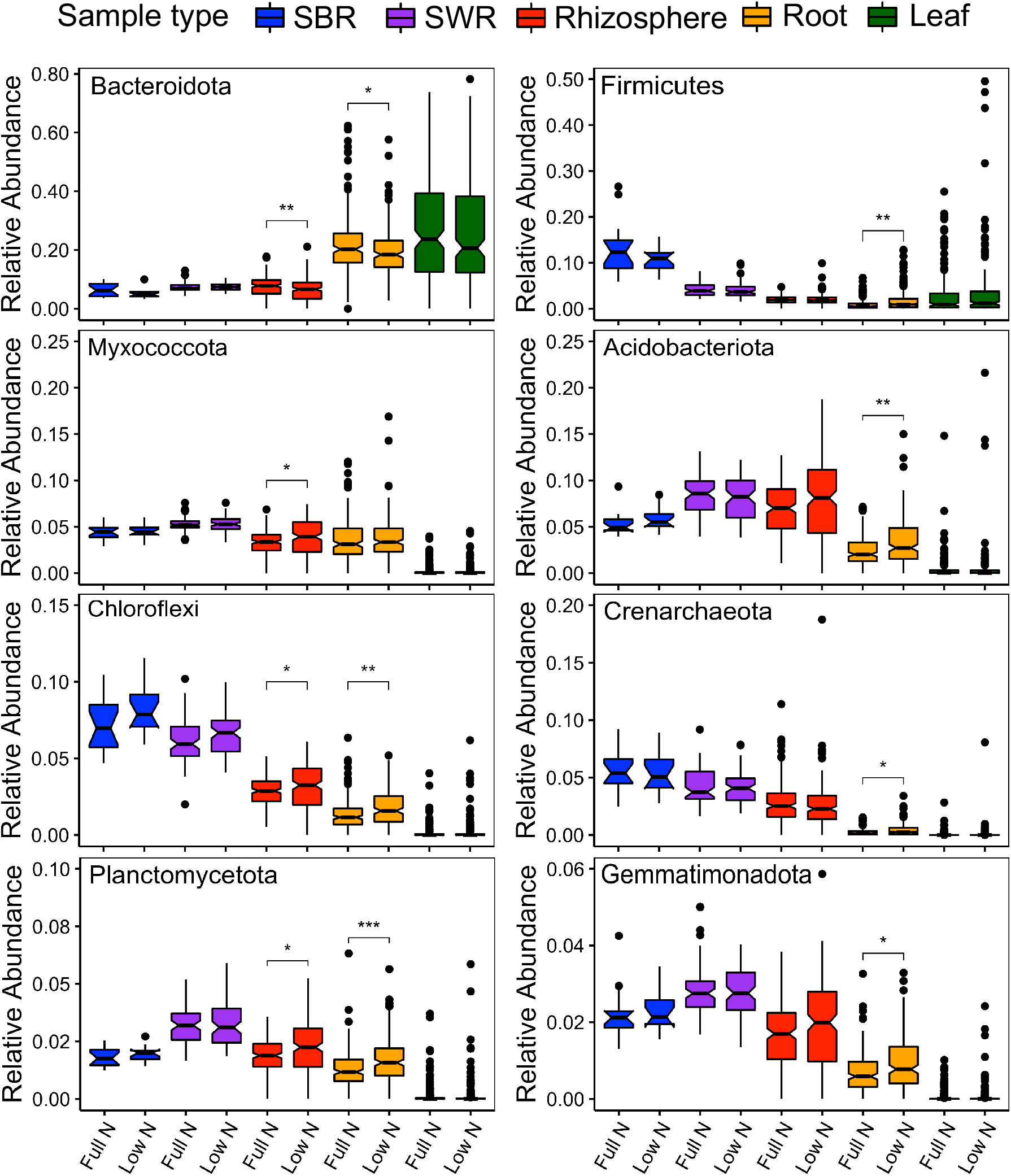
Relative abundance of bacterial phyla that were significantly impacted by N-stress in at least one compartment determined by LEfSE. *p ≤ 0.05, **p ≤ 0.01, ***p ≤ Abbreviation: SWR: soil within rows; SBR: soil between rows.

### The alpha diversity and composition of the rhizosphere bacterial community is associated with sorghum NUE

We sought to understand whether the shift in bacterial communities in the rhizosphere and root endosphere due to low-N were related to sorghum NUE. First, we correlated the changes in bacterial richness and diversity due to N stress with sorghum NUE expressed as the dry biomass ratio (Figure 1) across all sorghum genotypes (Figure 5A,B). The low-N-induced changes in the bacterial richness and diversity low-N were derived as the ratios of Faith’s PD and Shannon diversity between low- and full-N conditions, respectively. We found a significant positive correlation between these ratios and sorghum NUE only in the rhizosphere and not in the root endosphere (Figure S2), indicating that the genotypes with higher NUE had greater bacterial richness and diversity in the rhizosphere under low-N compared with full-N. This trend was stronger for the energy and grain sorghum (Figure 5C,D), but it is important to note that only two grain sorghum genotypes were included in this study.

**Figure 5.**
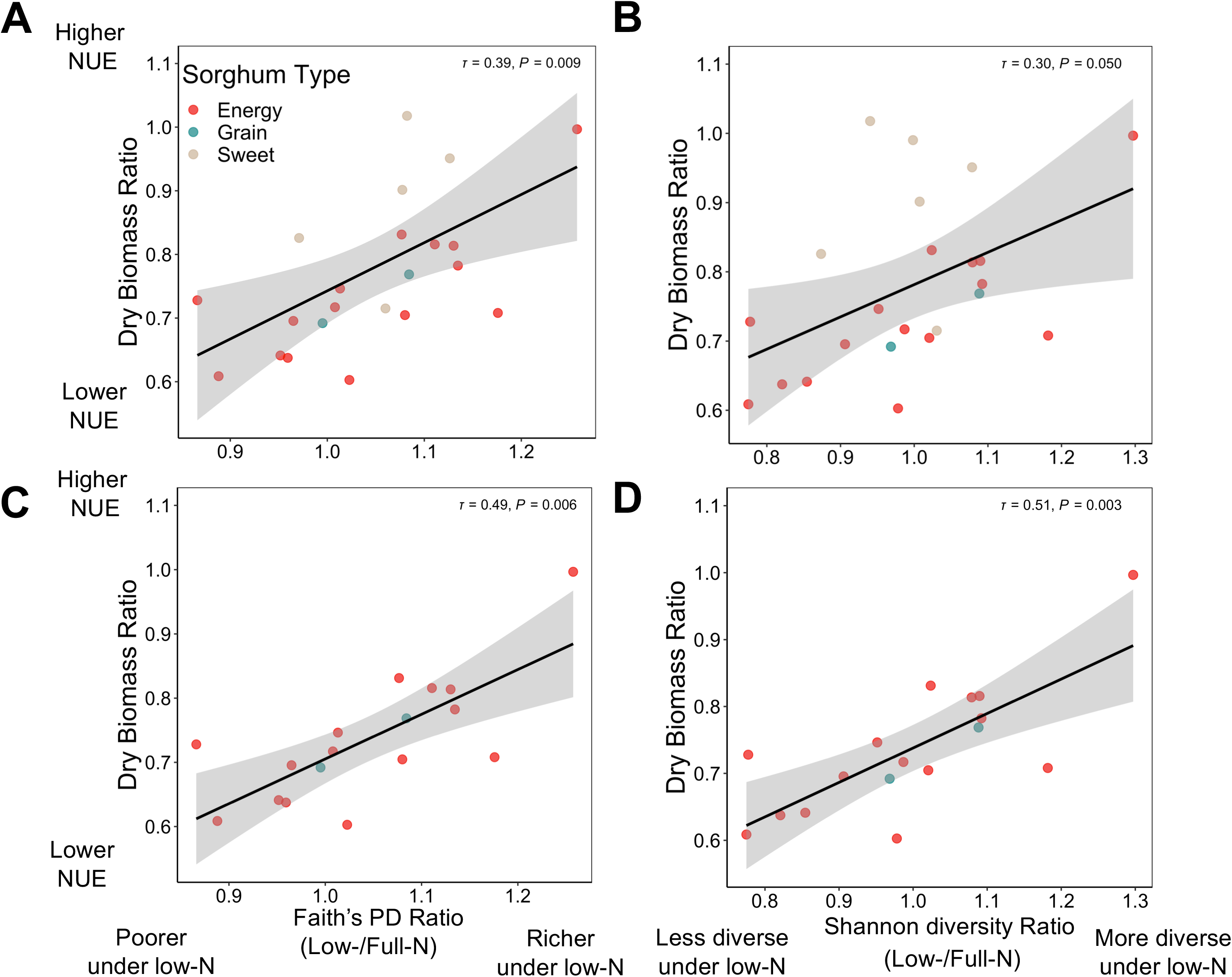
The relationship between bacterial alpha diversities in rhizosphere and sorghum NUE. (A,C) The correlation between Faith’s phylogenetic distance, (B,D) Shannon diversity ratio (low-N/full-N) with sorghum NUE derived from dry biomass ratio with (top) and without sweet sorghum (bottom). Coefficient of correlation as Kendall’s *τ* and *p*-value of each model is denoted on the top right of each plot.

We then investigated the effect of sorghum NUE on individual bacterial taxa using correlation analysis. We selected the microbial taxa across all taxonomic ranks that were responsive to low-N as identified by LEfSE followed by correlating their relative abundance ratio between low- and full-N with sorghum NUE. We found that the abundance ratio of *Pseudomonas*, the most abundant bacterial genus in the rhizosphere, was negatively correlated with NUE across the energy and grain sorghum type, but not for sweet sorghum (Figure 6A). In other words, *Pseudomonas* was more abundant in the rhizosphere of the energy and grain sorghum genotypes with lower NUE while it was less abundant in the rhizosphere of the genotypes with higher NUE under low-N. Since *Pseudomonas* comprised an average of 27 % of the rhizosphere bacterial community of energy and grain sorghum, we hypothesized that lower abundance of this genus in the genotypes with higher NUE under low-N may be related to the high alpha diversity in the rhizosphere of these sorghum genotypes, and *vice versa* for the low NUE lines. This hypothesis was tested by correlating the relative abundance of *Pseudomonas* in the rhizosphere to the Faith’s PD and Shannon diversity (Figure S3A,B). Strong negative correlations were detected between the abundance of *Pseudomonas* and alpha diversity, suggesting the differential abundance pattern of *Pseudomonas* in response to low-N was one factor that led to the differentiation of the rhizosphere bacterial communities across sorghum genotypes with different NUE. The sweet sorghum genotypes also had a lower abundance of *Pseudomonas* (∼ 20 %) in the rhizosphere than the other two sorghum types which corresponded to higher bacterial richness and diversity in sweet sorghum’s rhizosphere. Sweet sorghum genotypes tested also on average had higher NUE than the grain or energy types.Besides *Pseudomonas*, we identified other bacterial taxa that were differentially enriched or depleted in the rhizosphere of the genotypes with higher or lower NUE. An increased abundance of the phyla of Chloroflexi, Planctomycetota, Verrucomicrobiota, Actinobacteriota, Acidobacteriota, and the order Rhizobiales under low-N was also observed in the genotypes with higher NUE (Figure 6B-G).

**Figure 6.**
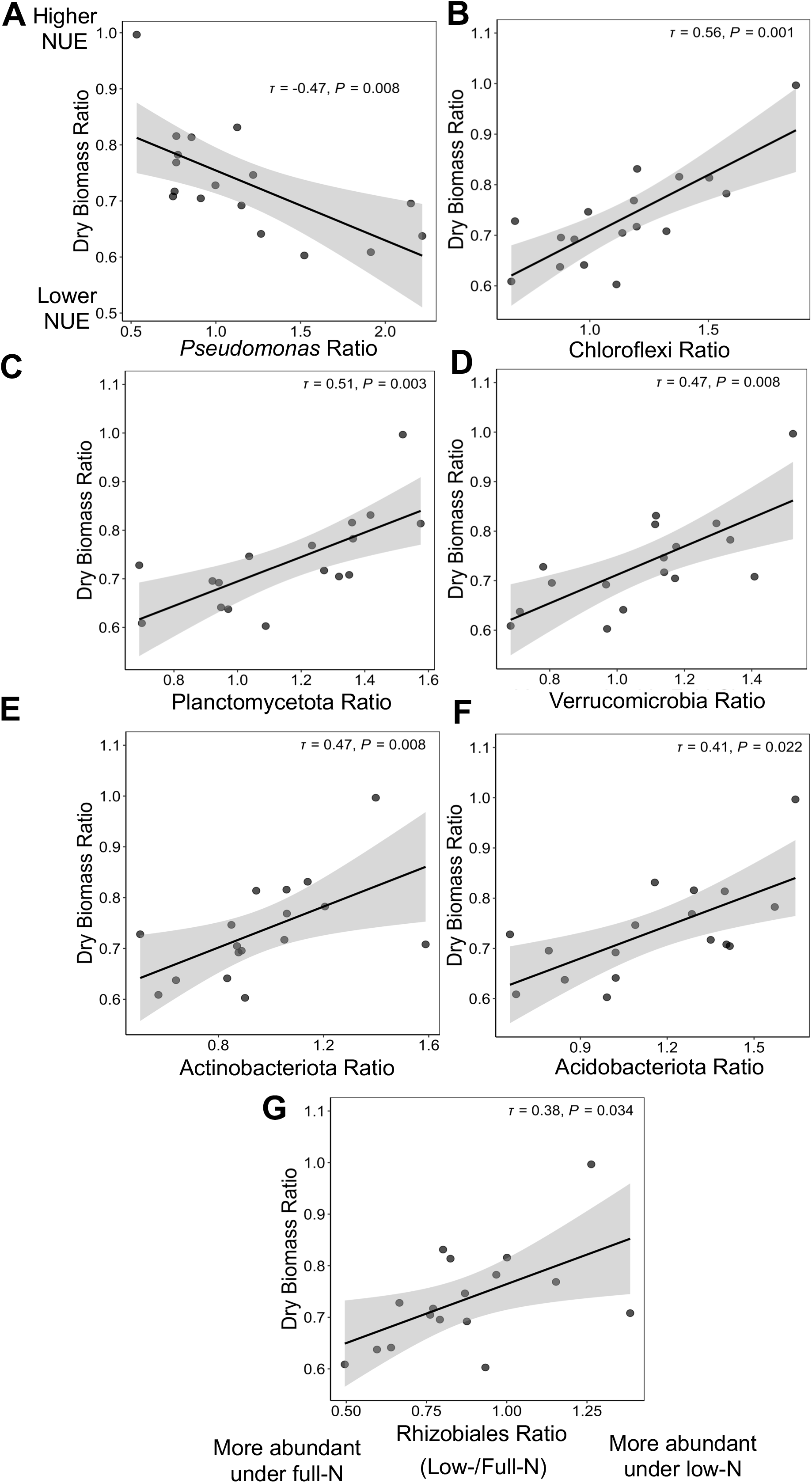
Bacterial taxa affected by sorghum NUE. Correlation of the differential abundance ratios of bacterial taxa under low-N/full-N versus sorghum NUE expressed as the biomass ratios under low-N/full-N (excluding sweet sorghum) of (A) Pseudomonas, (B) Chloflexi, (C) Planctomyctota, (D) Verrucomicrobiota, (E) Actinobacteriota, (F) Acidobacterriota, and (G) Rhizobiales. Coefficient of correlation as Kendall’s *τ* and p-value of each model is denoted on the top right of each plot.

### Sorghum root metabolite composition are affected by N availability and sorghum type and NUE

To determine whether low-N and sorghum type affected root metabolism, untargeted GC-MS was deployed to characterize root metabolites. The metabolomic profiling was performed on 19 sorghum genotypes (Table 1) with five replicates for each genotype and treatment combination. A total of 459 metabolites were detected and annotations were assigned to 77. PCA was used to visualize the variation across samples and showed that the root metabolite profile of the N-stressed plants was clearly different from those that were grown in the full-N segments of the field (Figure 7A).

**Figure 7.**
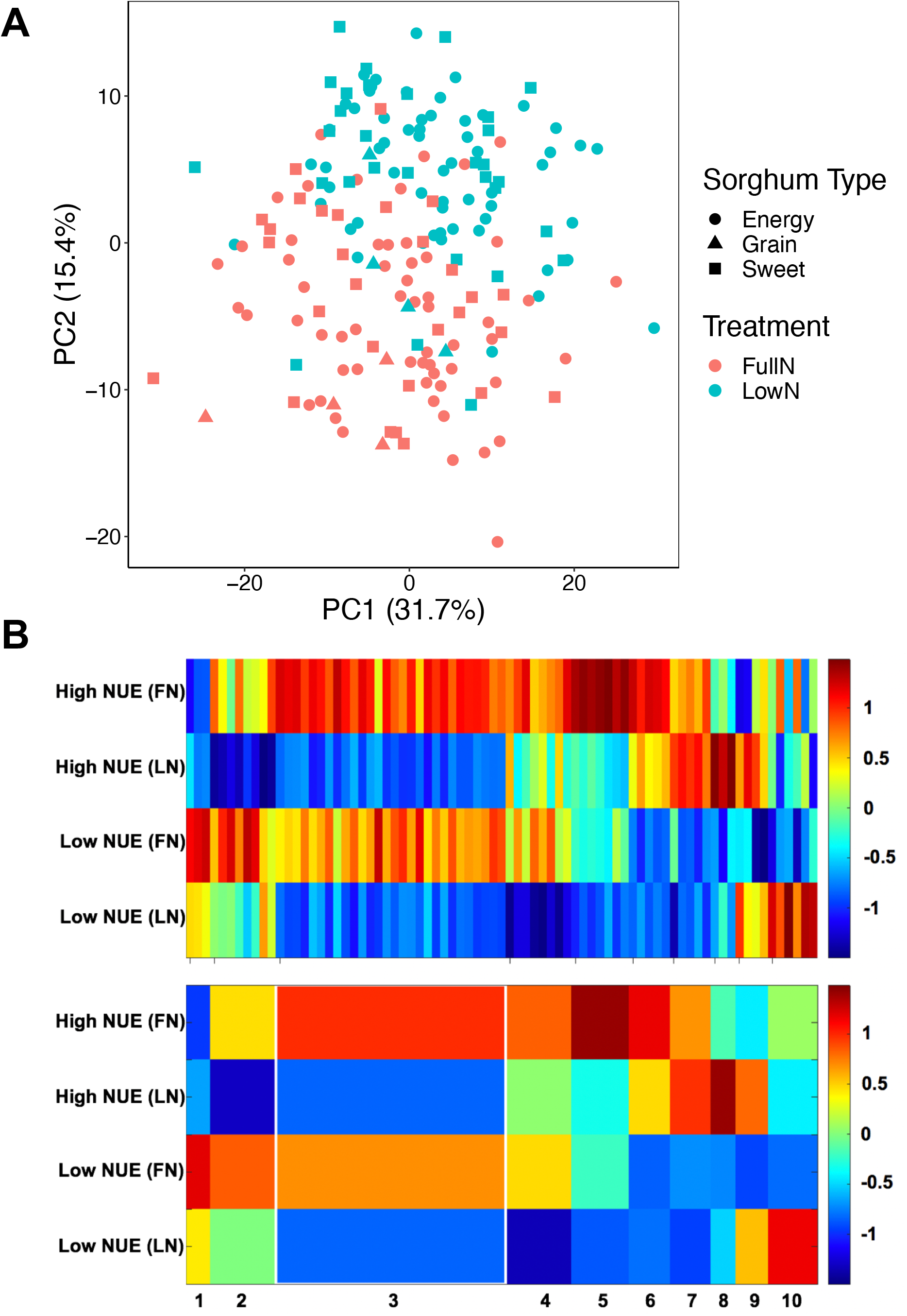
Root metabolomic profiles of sorghum. (A) PCA depicting the effects of N treatment and sorghum type on root metabolite composition. (B) 1D-SOM clustering and cluster assignment of sorghum root metabolites. The upper heatmap illustrates the relative signal intensity of each metabolite in high (biomass ratio > 0.78) and low NUE lines (biomass ratio < 0.78) under different N conditions. These metabolites were assigned to ten clusters at corresponding array position in the lower heatmap based on their intensity patterns. The color of each cluster represents the average intensity value among the metabolites in the same group. The width of each cluster corresponds to the number of metabolites assigned to the group. Abbreviation: FN: full-N; LN: low-N.

PERMANOVA confirmed that both low-N (*P* < 0.001) and sorghum type (*P* < 0.001) significantly affected root metabolite profile, but the effect from N stress was greater (*R*^*2*^ = 0.14) than that of sorghum type (*R*^*2*^ = 0.03).

A one-dimensional self-organizing map (1D-SOM) was used to cluster metabolites into ten clusters that showed similar patterns of systemic change (Fig 7B). (Supplemental dataset 1). The analysis revealed that a large number of metabolites decreased in intensity under N stress (clusters 2 - 5). The majority of the metabolites that decreased under low-N were amino acids (asparagine, glycine, leucine, lysine, phenylalanine, serine, and threonine) and organic acids (citric acid, succinic acid, lactic acid, and etc.). Despite showing decreased intensity under low-N, we found that metabolites in cluster 2 including trehalose and melibiose were more abundant in the low NUE lines. Conversely, cluster 8 and 9 featured metabolites that were enriched under low-N (Fig. 7B and Supplemental dataset 1) (ethanolamine, glucopyranose, galactose, organic acids as well as others) Strikingly, cluster 1 and clusters 6&7 clearly distinguished the metabolites that were differentially enriched in the high- and low-NUE lines, respectively. The metabolites that were enriched in low-NUE lines included shikimic acid while the metabolites enriched in high-NUE lines included tyrosine, fructose, and psicose (Figure 7B and Supplemental dataset 1). Cluster 10 represented the metabolites enriched in low-NUE lines specifically under N stress and these metabolites included hydroxylamine, pyridoxamine, phytol, itaconic acid, fumaric acid, and protocatechuic acid.

### Sorghum root metabolites shape the rhizosphere bacterial community

To determine whether the shifts in the rhizosphere bacterial community were induced by sorghum root metabolites, group sparse canonical correlation analysis (GSCCA) was performed to identify the correlation structure between metabolites and ASVs. Among the features selected by GSCCA, we identified several metabolites that were correlated with *Pseudomonas* ASVs in the rhizosphere. Despite the fact that 316 ASVs were assigned to *Pseudomonas,* the majority of the *Pseudomonas* reads (∼91%) was mapped to ten ASVs and eight out of these ten ASVs were enriched under low-N conditions. Interestingly, we found that trehalose was positively correlated with the dominant *Pseudomonas* ASVs (ASV2, 3, 9, 10, 16; *r* ∼ 0.27) that were specifically enriched under low-N. We then consolidated all the *Pseudomonas* ASVs and analyzed the correlation between *Pseudomonas* and root metabolites. We found that the ratio of *Pseudomonas* between low- and full-N exhibited a significant positive correlation with the ratio of shikimic acid between low- and full-N for all sorghum genotypes tested (Fig. 8A).

**Figure 8.**
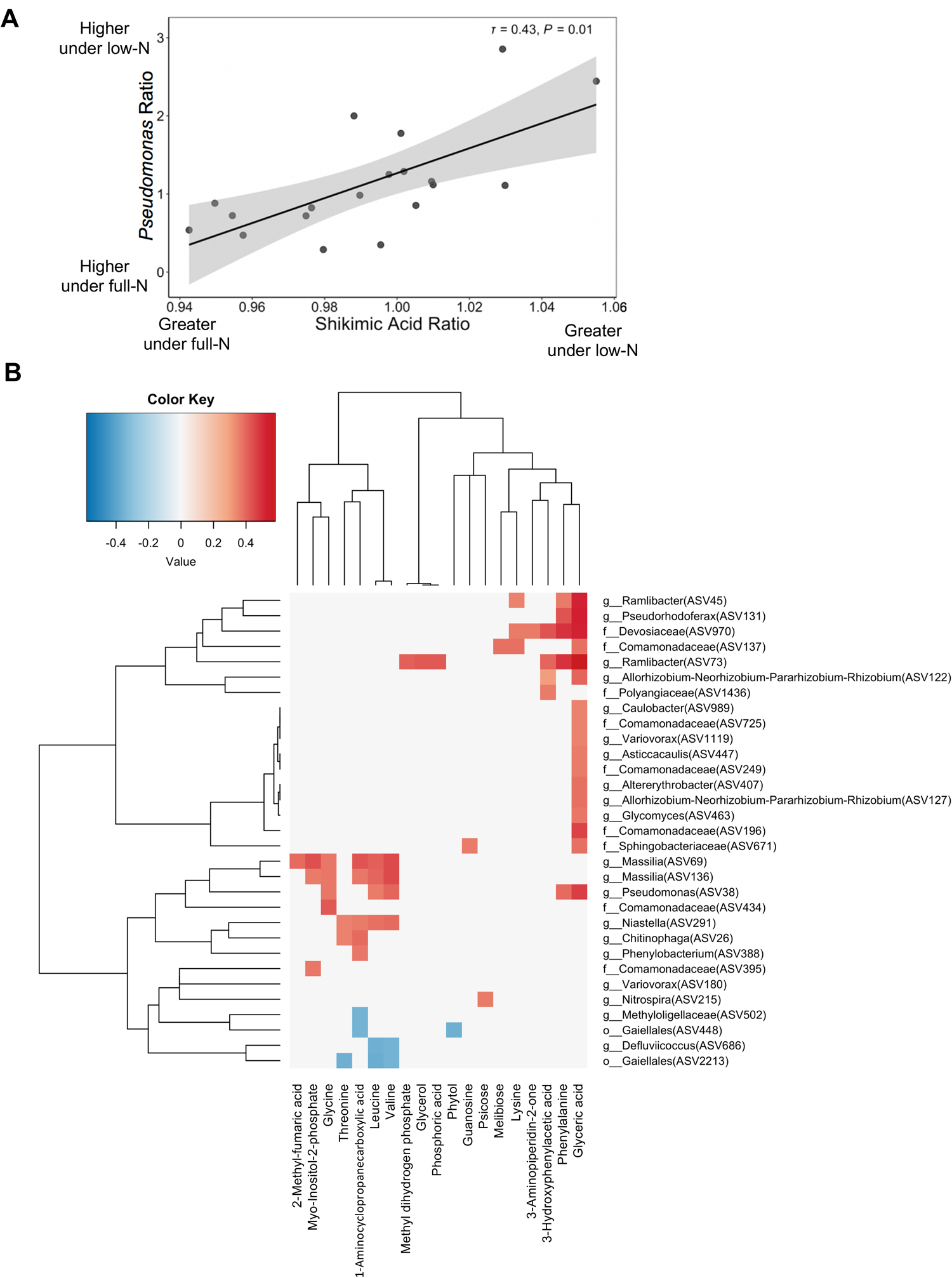
Relationship between root metabolites and rhizosphere bacteria. (A) Correlation between *Pseudomonas* ratio (low-N/full-N) and shikimic acid ratio (low-N/full-N). (B) Heatmap showing metabolites and ASVs identified by GSCCA with pairwise correlations were greater than 0.30.

In addition to *Pseudomonas*, GSCCA identified a number of ASV and metabolite pairs that were correlated (Figure 8B, Supplemental dataset 2). In particular, we found that psicose which was enriched in the high-NUE lines exhibited negative correlations with mostly the ASVs within the phyla of Actinobacteria and Proteobacteria while positively correlated with the members of Bacteroidota, Acidobacteriota, Methylomirabilota, Chloroflexi, Gemmatimonadota, and numerous potential nitrifying taxa including *Nitrospira, Ellin6067, Rokubacteriales*, and *NB1-J* (Figure 8B, Supplemental dataset 2). On the other hand, phytol which was enriched in low-NUE lines also covaried with many members of Actinobacteria and Proteobacteria, but the impact was specific to lower taxonomic rank. For example, phytol was positively correlated with *Streptomyces* while negatively correlated with Gaiellales, *Mycobacterium, Aeromicrobium, Nocardioides*, and *Solirubrobacter* even though they are all members of Actinobacteria. GSCCA also identified a number of ASVs that were correlated with organic acids, amino acids, and sugars (Figure 8B, Supplemental dataset 2).

## Discussion

The communication between soil microbial communities and plant root metabolites and exudates is complex and varies depending on plant species, environmental conditions, plant developmental stage as well as many soil physicochemical factors. It is well known that root metabolites such as nodulation (NOD) factors stimulate the beneficial interaction between nitrogen fixing legumes and Rhizobia under conditions of low N supply. Under conditions of biotic stress the idea of a “cry for help” has been discussed as a way in which plants deploy metabolite signals to recruit beneficial microbes to help alleviate both abiotic and biotic stresses (Dicke & Baldwin, 2010; Liu et al., 2021; Rizaludin et al., 2021; Rolfe et al., 2019). While much of the research enumerating examples of the “cry for help” hypothesis has measured root exudates – root metabolites also provide an important factor in shaping microbial communities (Bourceret et al., 2022; Csorba et al., 2022; DeWolf et al., 2022).

Improving NUE in cereal crops is of critical importance to agriculture worldwide. Conventional breeding and genetic engineering are the main approaches currently used to enhance crop NUE (Han et al., 2015). The yield enhancement resulting from agronomic approaches where increasing amounts of N fertilizer are used may no longer be a feasible approach to improve crop yield because of the environmental externalities caused by fertilizer usage. Breeding for increased NUE has also been very challenging because it has been difficult to identify genes responsible for NUE due to the significant variability in field testing approaches and the likelihood that this is a quantitative trait (Han et al., 2015). New methods to improve crop NUE by manipulating plant microbiomes may be a solution that provides a more sustainable alternative since new evidence shows that crop NUE may in part be associated with the plant belowground microbiome (Zhang et al., 2019).

### NUE characteristics in sorghum

The genetic variability of sorghum contributes to significant variation in the NUE across different sorghum genotypes (Bollam et al., 2021). In this study, we compared the growth of 24 diverse sorghum lines under two N regimes and identified the genotypes that exhibited higher NUE when N was limited. Consistent with other studies (Chai et al., 2021; Gelli et al., 2014), we found that sweet sorghum genotypes generally have higher NUE as compared to energy and grain sorghum. PI 297155 was one of the genotypes with highest NUE identified in this study, but it was also determined to have the lowest NUE in our previous greenhouse study (Chai et al., 2021). This discrepancy in sorghum NUE between greenhouse and field conditions is not surprising because crop NUE could vary with many factors including N fertilization rates (Barraclough et al., 2010) and soil characteristics (e.g., pH, texture) (Ichami et al., 2019), highlighting the importance of selecting for crop NUE in different environments. Overall, our study highlights the large degree of NUE variability across diverse sorghum genotypes and provides insight on the potential candidates to be used in breeding of high NUE sorghum lines.

### Role of Microbiome on Influencing Sorghum NUE

Microbial richness and diversity are frequently demonstrated to be positively correlated with plant vigor (Berg et al., 2017; Chen et al., 2020; Jaiswal et al., 2017; Zhang et al., 2019). Yet, our knowledge of the mechanisms underpinning the positive effects of microbial diversity on plant health are limited to the perspective that higher microbial diversity helps buffer against the perturbation of certain microbial species that are usually harmful to plants (Berg et al., 2017). Our finding that increased microbial richness and diversity in the rhizosphere are positively correlated with plant performance under low-N conditions, provides evidence that increased bacterial diversity may contribute to mitigating sorghum N-stress. This correlation is also indicative of the large variation in responses to low-N among the range of sorghum lines studied here that differ in NUE. In agreement with our result, greater bacterial diversity has also been found in the root endosphere of *indica* rice varieties which exhibits superior NUE as compared to the *japonica* lines (Zhang et al., 2019). In the rice study the increased bacterial diversity was attributed to the recruitment of a large proportion of N-cycle related bacteria of *indica* varieties which may have enhanced the plant N availability. In another study on tree species when N became limited, plant root metabolites were shown to stimulate microbial decomposition of soil organic matter to increase N mineralization for plant uptake (Dijkstra et al., 2013; Dijkstra et al., 2009).Our findings and those of others suggest that high NUE in plants may in part be linked to the ability to modulate the belowground microbial communities to ultimately improve N-uptake and availability. The positive correlation between microbial diversity and plant NUE may be due to enhanced complementary interactions (Loreau, 2000) and improved overall metabolic capabilities of rhizosphere microbial community under low-N. This hypothesis warrants further testing given the importance of increasing NUE in modern agroecosystems. A functional consequence of the increased diversity may be to boost soil organic matter decomposition and other aspects of N-cycling with positive impacts on the overall productivity of the whole plant-microbe holobiont.

We further revealed that the increased bacterial diversity and richness in the rhizosphere of high NUE genotypes under low-N were associated with the decreased abundance from *Pseudomonas* and increased abundance of the members from Chloflexi, Planctomycetota, Verrucomicrobiota, Actinobacteriota, Acidobacteriota, and *Rhizobiales. Pseudomonas* was the most abundant genus identified in the rhizosphere in our study. Strikingly, *Pseudomonas* proliferation particularly during the early host vegetative stage has also been documented in various independent studies on sorghum (Chiniquy et al., 2021; Lopes et al., 2021; Sheflin et al., 2019) and maize (Walters et al., 2018). Although the cause of the *Pseudomonas* proliferation is unclear, *Pseudomonas* is a copiotroph specialized in utilizing plant root metabolites (Chiniquy et al., 2021).

Therefore, we hypothesize that the lower abundance of *Pseudomonas* in the high NUE genotypes may be driven by reduced access to labile carbon from plant roots in response to N-stress, which facilitated the enrichment of the oligotrophic phyla, Chloflexi, Planctomycetota, Verrucomicrobiota, and Acidobacteria (Ho et al., 2017). The enrichment of the order *Rhizobiales* in the rhizosphere may also contribute to plant N uptake as this order contains many N-fixing genera (Erlacher et al., 2015). The association between the relative abundance of these taxa and host NUE was stronger in energy and grain, but not in sweet sorghum genotypes which exhibited greater NUE. This suggested that the effect of rhizosphere bacterial communities on sorghum NUE may be related to the sorghum types and the degree of NUE in various genotypes.

### The modulation of root metabolites under low-N

N-availability was a main factor impacting the root metabolite composition in our study, with the decrease in many amino acids being the most pronounced change in N-stressed roots. This finding is consistent with other studies (Sheflin et al., 2019; von Wirén et al., 2000) and subsequently led to the reduction of several hydroxycinnamates and hydroxybenzoates derived from phenylpropanoid pathway. Many of these compounds (*p*-coumaric acid, vanillic acid, protocatechuic acid, *p*-benzoic acid, and ferulic acid) are integral constituents of cell wall and involved in plant defense against pathogens (Vogt, 2010). Therefore, the reduction in these compounds under low-N may have accounted for the increase in bacterial richness demonstrated in the rhizosphere and root endosphere.

We also demonstrated that certain root metabolites were differentially enriched or depleted in sorghum lines with different NUE, suggesting that there may be associations between root metabolism and sorghum NUE. For instance, we found that, psicose, a rare sugar found in plants was differentially enriched in the root of high-NUE lines regardless of the N treatment. Psicose, also known as allulose, has been shown to induce rice resistance against bacterial blight through upregulation of many defense-related genes (Kano et al., 2010). Psicose was also found to be positively correlated with several potential nitrifying genera including *Nitrospira* (Lücker et al., 2010), *Ellin6067* (Wang et al., 2021), *Rokubacteriales* (Becraft et al., 2017), and *NB1-J* (de Voogd et al., 2015). Our findings that psicose was enriched in the high-NUE lines and positively correlated with potential nitrifying bacteria may indicate that a larger supply of nitrate was available to the higher NUE lines. While the scope of our study was insufficient to test the function of psicose accumulation in high-NUE lines, we also noticed that it might have had a suppressive effect on the copiotrophic phyla because of the negative correlation between psicose levels and numerous members of Actinobacteria and Proteobacteria (Leff et al., 2015).

In contrast to the positive association between psicose and NUE, phytol accumulation was associated with low-NUE lines under N stress. Phytol is a diterpene constituent of chlorophyll and the accumulation of phytol is often associated with chlorophyll degradation induced by stresses such as N deprivation (Gutbrod et al., 2021; vom Dorp et al., 2015). Therefore, greater phytol accumulation in roots in the low-NUE lines further shows that N stress was more severe than that of the high-NUE genotypes. We also showed that the shikimic acid and trehalose levels in roots were positively correlated with the *Pseudomonas* ASVs in the rhizosphere of low-NUE lines under N stress. In addition, both the work by Sheflin et al. 2019 and our study found that the concentration of shikimic acid in the sorghum roots was correlated to the differential abundance of *Pseudomonas* in the rhizosphere. Shikimic acid is a precursor of aromatic amino acids including phenylalanine, which is required for the synthesis of a plant defense hormone, salicylic acid. Although root salicylic acid was not measured, our results provide further evidence that the differential abundance of *Pseudomonas* in this study may be attributed to the varying defense or stress responses in the sorghum lines that differed in NUE under low N conditions (Sheflin et al., 2019). Trehalose has been shown to have integral roles in alleviating plant abiotic stresses such as drought and salinity (Fernandez et al., 2010). While little is known about the role of trehalose, in our study it was related to reduced microbial diversity and lower NUE in sorghum. In contrast under nitrogen deficiency an association between cereal species and tolerance to low nitrogen conditions was linked to trehalose (Sun et al., 2022) but no comparisons were done between genotypes of the same species as in our study. Taken together, our findings highlight the potential crosstalk between host root metabolites and the rhizosphere bacterial communities that may be important for plant performance under low-N conditions.

## Conclusion

The objectives of this study were to more fully elucidate how sorghum genotype, root associated bacteria and root metabolites may be related to nitrogen use efficiency in sorghum. The findings from our study demonstrate that there are associations between the sorghum root metabolome and rhizosphere bacterial communities that are associated with sorghum NUE across a range of genetically diverse genotypes. We found that sorghum bacterial communities were impacted by sorghum type (grain, sweet and energy). N-stress was also an important factor that led to change in the composition of bacterial communities in the rhizosphere and root endosphere. Although the shifts in root metabolite composition were mainly driven by N availability, sorghum type did influence root metabolite composition. Sorghum NUE was positively correlated with the richness and diversity in the bacterial community in the rhizosphere, which was driven by the differential abundance of the dominant *Pseudomonas* genera. We also showed that the changes in root metabolites due to N-stress were correlated with the shifts in the rhizosphere bacterial community composition. Taken together, our results highlight the importance of considering the vast sorghum genetic variation in both plant response to changes in soil fertility which led to certain fingerprints of root metabolome and root-associated bacterial community composition that were related to the differences in sorghum NUE. Future experiments will be needed to test whether specific microbes and metabolites identified in this study actually contribute to sorghum NUE.

## Acknowledgements

The work was supported by a grant from the US Department of Energy award DE_SC0014395. Parts of the work were also (proposal: 10.46936/10.25585/60001066) conducted by the U.S. Department of Energy Joint Genome Institute (https://ror.org/04xm1d337), a DOE Office of Science User Facility, is supported by the Office of Science of the U.S. Department of Energy operated under Contract No. DE-AC02-05CH11231. We thanks Stephane Futrell, Bert Devilbiss, John Rajewski, Glenn Slater for expert technical assistance and Stephen Kresovich, Ismail Dweikat and Bill Rooney for providing seed used in these experiments.

## Figure legend

**Figure S1.**
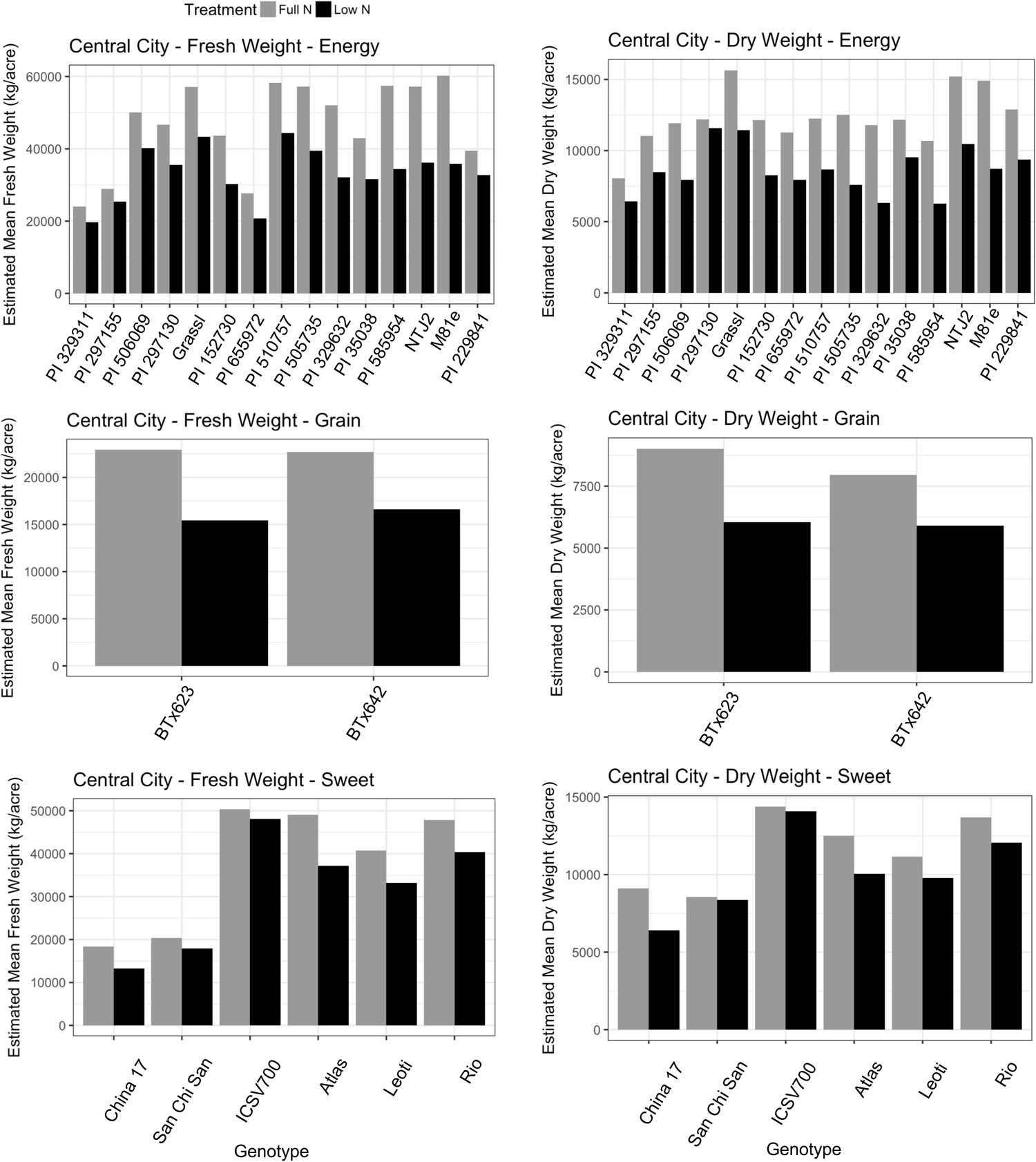
Biomass of 23 diverse sorghum genotypes. Fresh (left) and dry weight (right) of each sorghum genotype under full- and low-N.

**Figure S2.**
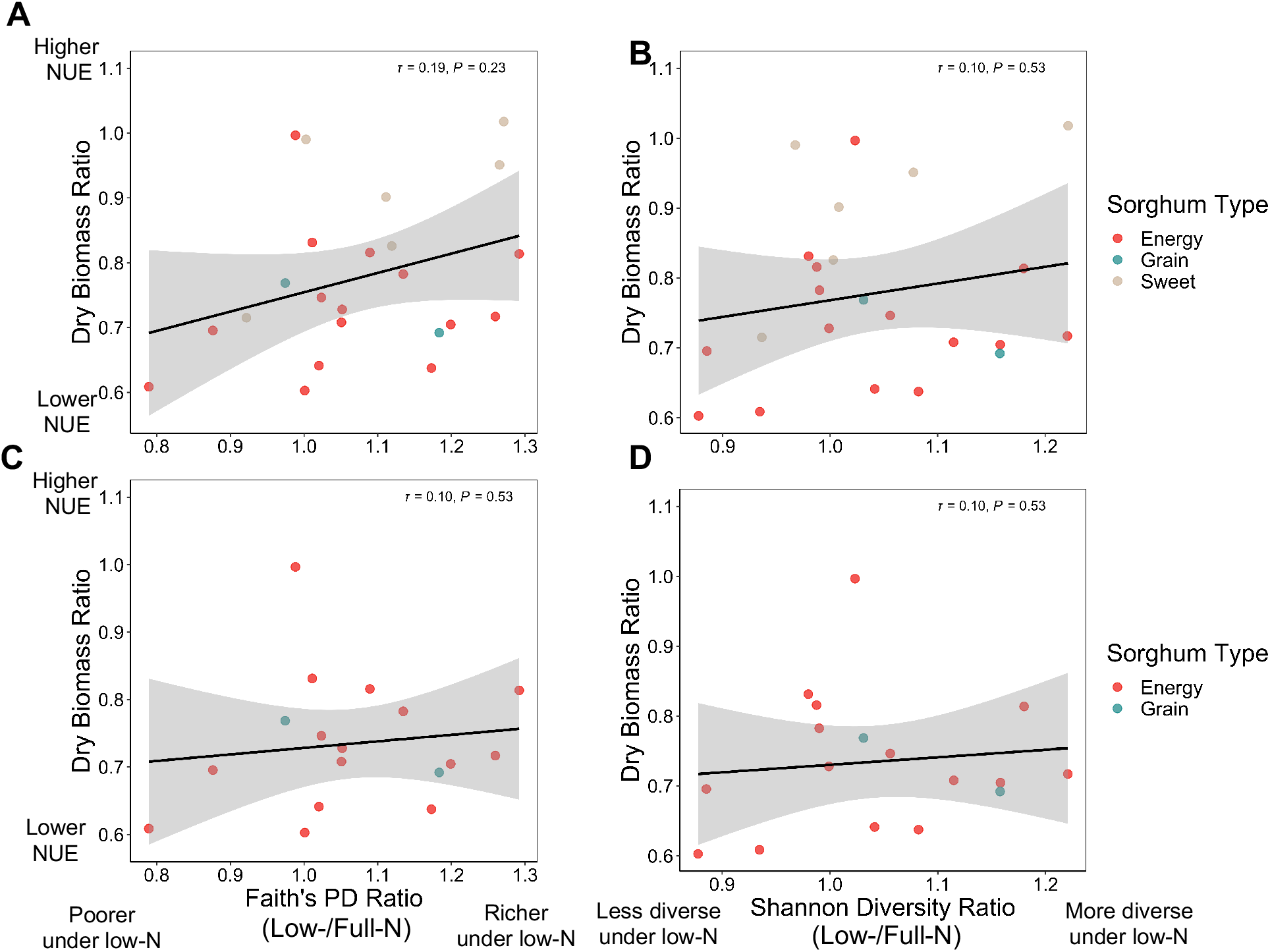
The relationship between bacterial alpha diversities in root endosphere and sorghum NUE. (A,C) The correlation between Faith’s phylogentic distance, (B,D) Shannon diversity ratio (low-N/full-N) with sorghum NUE derived from dry biomass ratio with (top) and without sweet sorghum (bottom). Coefficient of correlation as Kendall’s *τ* and *p*-value of each model is denoted on the top right of each plot.

**Figure S3.**
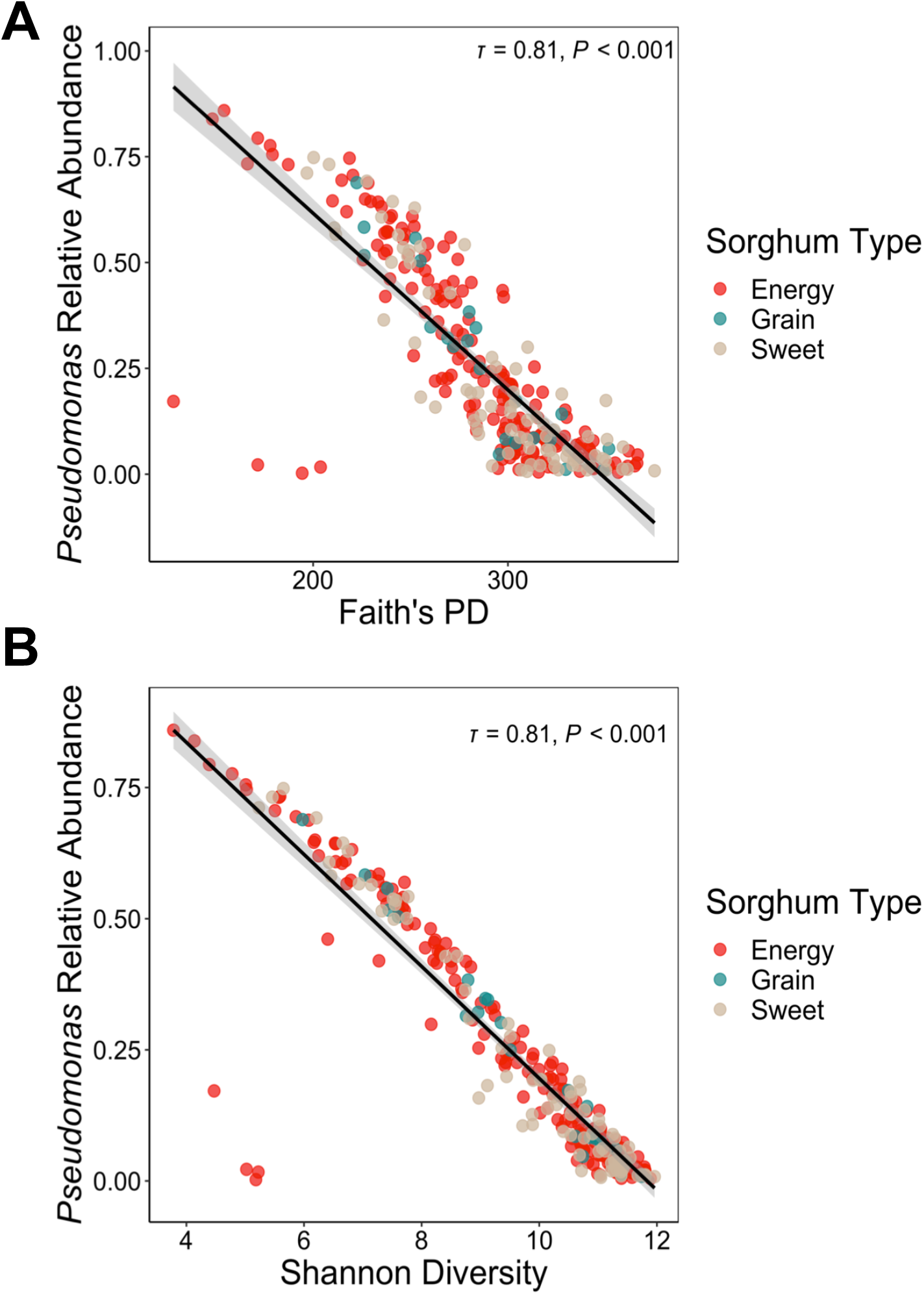
Relationship between alpha diversities and *Pseudomonas* abundance. Correlation between (A) Faith’s phylogenetic distance, (B) Shannon diversity with *Pseudomonas* abundance.

